# Autophagy controls lipid droplet formation by fine-tuning NCoR1 levels

**DOI:** 10.1101/722686

**Authors:** Shun-saku Takahashi, Yu-Shin Sou, Tetsuya Saito, Akiko Kuma, Takayuki Yabe, Masato Koike, Shuji Terai, Noboru Mizushima, Satoshi Waguri, Masaaki Komatsu

**Affiliations:** Division of Gastroenterology and Hepatology, Niigata University Graduate School of Medical and Dental Sciences, Chuo-ku, Niigata 951-8510, Japan; Department of Cell Biology and Neuroscience, Juntendo University Graduate School of Medicine, Bunkyo-ku, Tokyo 113-8421, Japan; Department of Physiology, Juntendo University Graduate School of Medicine, Bunkyo-ku, Tokyo 113-8421, Japan; Department of Biochemistry and Molecular Biology, Graduate School and Faculty of Medicine, the University of Tokyo, Bunkyo-ku, Tokyo 113-0033, Japan; Department of Physiology and Cell Biology, Tokyo Medical and Dental University, Tokyo 113-8519, Japan; Department of Anatomy and Histology, Fukushima Medical University School of Medicine, Hikarigaoka, Fukushima 960-1295, Japan

**Keywords:** autophagy, NCoR1, LXRα, lipid droplet

## Abstract

Lipid droplets (LDs) are dynamic organelles that store neutral lipids during times of energy excess, such as following a meal. LDs serve as an energy reservoir during fasting and have a buffering capacity that prevents lipotoxicity. Autophagy and the autophagic machinery have been proposed to play a role in LD biogenesis but the underlying molecular mechanism remains unclear. Here, we show that when nuclear receptor co-repressor 1 (NCoR1), which inhibits the transactivation of nuclear receptors, accumulates due to autophagy suppression, LD biogenesis is blocked. Ablation of *ATG7*, a gene essential for autophagy, suppressed the expression of gene targets of liver X receptor α (LXRα), a nuclear receptor responsible for fatty acid and triglyceride synthesis in an NCoR1-dependent manner. LD biogenesis in response to fasting and after hepatectomy was hampered by the suppression of autophagy. These results indicate that autophagy controls physiological hepatosteatosis by fine-tuning NCoR1 protein levels.

## Introduction

Lipid droplets (LDs) are neutral lipid storage organelles that provide fatty acids (FAs) for energy production during periods of nutrient deprivation. These organelles, which emerge from the endoplasmic reticulum (ER), also have a lipid buffering capacity that helps prevent lipotoxicity (Henne, Reese, & Goodman, 2018; Olzmann & Carvalho, 2019). Enzymes involved in triacylglycerol (TG) synthesis, such as diacylglycerol O-acyltransferase (DGAT), deposit neutral lipids in between the leaflets of the ER bilayer where neutral lipids demix and coalesce to form a structure called an oil lens. Thereafter, seipin and other lipid droplet biogenesis factors facilitate the growth of nascent LDs from this lens. LDs bud from the ER and grow through either fusion or local lipid synthesis (Henne et al., 2018; Olzmann & Carvalho, 2019).

Apart from the selective degradation of LDs (lipophagy) (Olzmann & Carvalho, 2019; Singh et al., 2009), there is growing evidence that autophagy or some element of the autophagic machinery plays an important role in LD biogenesis. First, there have been several independent observations of a reduction in lipid droplet number in knockout mice lacking autophagic components specifically in their hepatocytes (Kim et al., 2013; Kwanten et al., 2016; Ma et al., 2013; Shibata et al., 2010; Takagi et al., 2016). Second, the autophagic machinery participates in lipid droplet formation in hepatocytes and cardiomyocytes (Shibata et al., 2009; Shibata et al., 2010), and deletion of autophagy-related genes such as *Atg5* and *Atg7* in the mouse liver decreases the level of triglycerides in the liver (Shibata et al., 2009) and impairs ketogenesis (Saito et al., 2019; Takagi et al., 2016). Third, the loss of Fip200, an autophagy initiation factor, in mouse livers causes inactivation of nuclear receptors, liver X receptor α (LXRα), and peroxisome proliferator-activated receptor α (PPARα). These receptors play important roles in FA synthesis and oxidation, respectively (Ma et al., 2013). Therefore, their inactivation blocks liver steatosis under physiological fasting and high-fat diet conditions (Ma et al., 2013). Fourth, the supply of lipids provided through autophagy is required to replenish triglycerides in lipid droplets (Rambold, Cohen, & Lippincott-Schwartz, 2015), which provide molecules for fatty acid oxidation. Fifth, the biogenesis of LD from FA supplied by starvation-induced autophagy prevents the lipotoxic effects of acylcarnitine (Nguyen et al., 2017), which disrupts mitochondrial membrane potential and mitochondrial function. However, whether autophagy participates in LD biogenesis directly and which step(s) within the process of LD biogenesis is affected by autophagy both remain unclear.

In this Short Report, we show that autophagy regulates FA and TG synthesis at the transcriptional level by fine-tuning the levels of nuclear receptor co-repressor 1 (NCoR1), a negative regulator of nuclear receptors, including LXRα, and that defective autophagy impairs LD biogenesis both under fasting conditions and following hepatectomy.

## Results and discussion

### Impairment of fasting-induced hepatosteatosis in liver-specific *Atg7*-knockout mice

NCoR1 is an autophagy-specific substrate (Iershov et al., 2019; Saito et al., 2019) and serves as a scaffold that facilitates the interaction of several docking proteins to fine-tune the transactivation of nuclear receptors such as LXRα and PPARα (Mottis, Mouchiroud, & Auwerx, 2013; Perissi, Jepsen, Glass, & Rosenfeld, 2010). The interaction of NCoR1 with nuclear receptors and histone deacetylases is vital for nuclear receptor–mediated downregulation of gene expression. Interestingly, LXRα and PPARα, both of which are negatively regulated by NCoR1, play opposing roles in lipid metabolism. Specifically, LXRα serves anabolic roles (FA and TG syntheses) while PPARα serves a catabolic role (β-oxidation). To determine whether NCoR1 accumulation due to autophagy suppression has an impact on LD biogenesis, we utilized hepatocyte-specific *Atg7*-knockout mice, *Atg7*^*f/f*^;Alb-*Cre* mice. The conversion of LC3-I to LC3-II was completely inhibited by the loss of *Atg7* (Fig. 1A), and p62/SQSTM1 (hereafter referred to p62), another autophagy-specific substrate, accumulated in mutant livers (Fig. 1A), implying that autophagy was impaired. In agreement with previous reports (Iershov et al., 2019; Saito et al., 2019), we found that NCoR1 accumulates in both the nuclear and cytoplasmic fractions prepared from livers of *Atg7*^*f/f*^;Alb-*Cre* mice (Fig. 1A). Fasting decreased NCoR1 in both fractions from mutant livers, but levels of this protein were still higher than in control livers (Fig. 1A).

**Figure 1.**
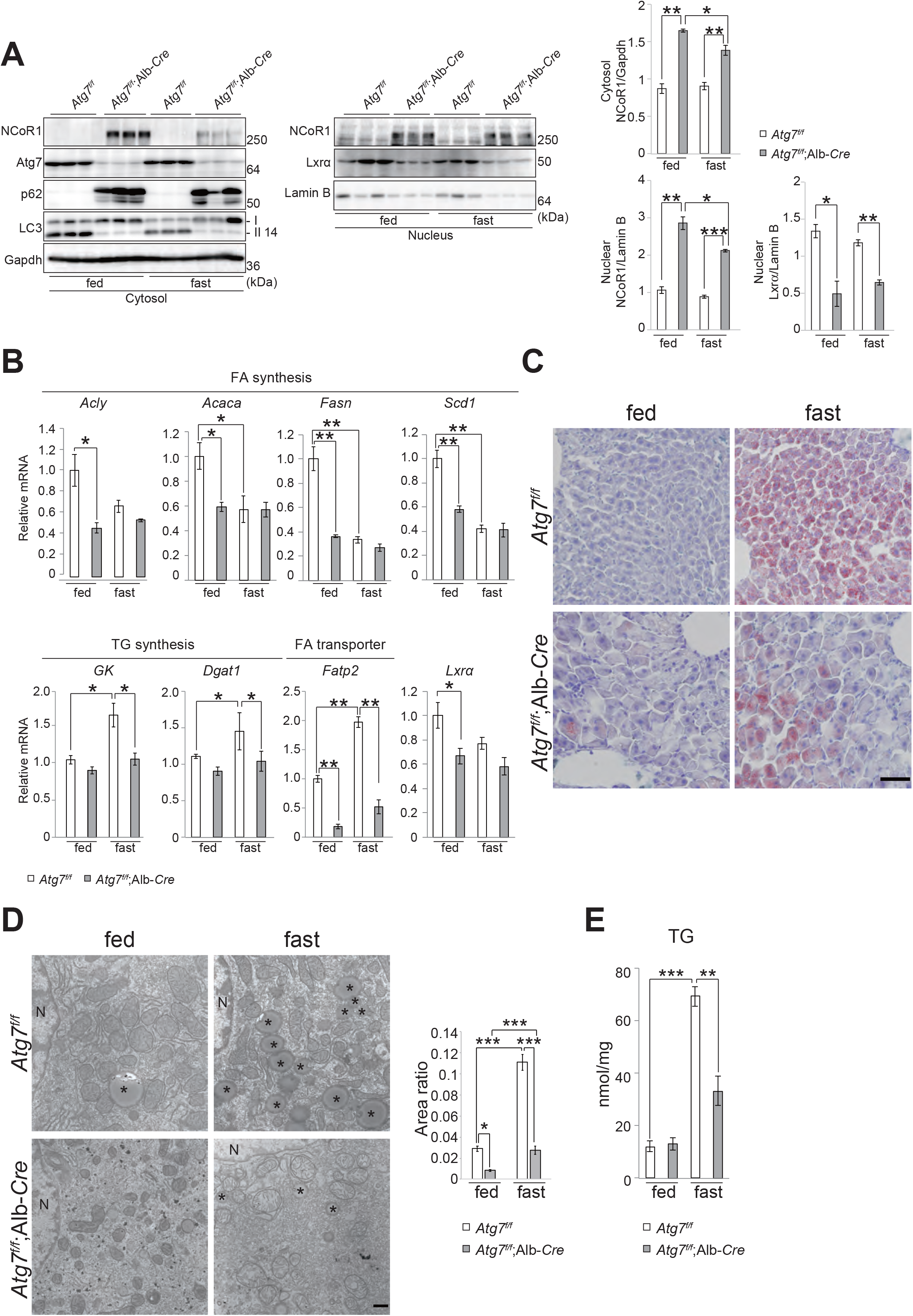
(**A**) LXRα and NCoR1 levels in *Atg7*-deficient mouse livers. Total homogenate, as well as nuclear and cytoplasmic fractions, were prepared from the livers of five-week-old *Atg7*^*f/f*^ and *Atg7*^*f/f*^;Alb-*Cre* mice under both fed and fasting conditions. These were subjected to immunoblotting using the indicated antibodies. Bar graphs indicate the amounts of the indicated cytoplasmic and nuclear proteins measured by densitometry relative to Gapdh and Lamin B, respectively. Data are means ± s.e. \**p* < 0.05, \*\**p* < 0.01, and \*\*\**p* < 0.001 as determined by Welch’s *t*-test. (**B**) Gene expression of proteins related to FA and TG syntheses in *Atg7*-deficient livers. Total RNA was prepared from the livers of five-week-old *Atg7*^*f/f*^ (n = 4) and *Atg7*^*f/f*^;Alb-*Cre* (n = 4) mice under both fed and fasting conditions. Values were normalized against the amount of mRNA in the livers of *Atg7*^*f/f*^ mice under fed conditions. The experiments were performed three times. Data are means ± s.e. \**p* < 0.05, and \*\**p* < 0.01 as determined by Welch’s *t*-test. (**C**) Oil-red O staining. Cryo-sections were prepared from livers of five-week-old *Atg7*^*f/f*^ and *Atg7*^*f/f*^;Alb-*Cre* mice under both fed and fasting conditions, and subjected to Oil-red O staining. Bar: 50 μm. (**D**) Electron microscopy. Representative electron micrographs of hepatocytes from the same genotype mouse as in (C) are shown. Ratio of LD area was measured and plotted in the right graph. Data are means ± s.e. \**p* < 0.05, and \*\*\**p* < 0.001 as determined by Welch’s *t*-test. Asterisks: LD, N: nucleus, Bar: 500 nm. (**E**) Liver triglyceride (TG) in mice described in (B). Data are means ± s.e. \*\**p* < 0.01, and \*\*\**p* < 0.001 as determined by Welch’s *t*-test.

The expression of genes encoding enzymes involved in FA synthesis, including *ATP citrate lyase* (*Acly*), *Acetyl-CoA carboxylase* (*Acaca*), *Fatty acid synthase* (*Fasn*), and *Stearoyl-CoA desaturase* (*Scd1*), which is regulated by LXRα, was markedly suppressed in the livers of *Atg7*^*f/f*^;Alb-*Cre* mice under fed conditions (Fig. 1B). Under fasting conditions, transcript levels of enzymes related to FA synthesis in control livers decreased to a similar extent as those in mutant livers (Fig. 1B). Remarkably, while the genes encoding enzymes related to TG synthesis, such as *Glycerol kinase* (*Gk*) and *Diacylglycerol O-acyltransferase* (*Dgat1*) and a transporter of FAs, *Fatty acid transport protein 2* (*Fatp2*), were upregulated upon fasting, such induction was hardly observed in livers of *Atg7*^*f/f*^;Alb-*Cre* mice (Fig. 1B). The level of *Lxrα* mRNA was lower in mutant livers than in control livers (Fig. 1B), consistent with the idea that LXRα regulates its own expression (Li et al., 2002). Since the autophagic turnover of NCoR1 is necessary for effective β-oxidation in response to fasting (Iershov et al., 2019; Saito et al., 2019), these results suggest that under fasting conditions, both the catabolism (β-oxidation) and anabolism (TG synthesis) of FAs are primed by NCoR1 degradation. In fact, LDs detected by Oil-red O staining and electron microscopy showed that fasting-induced hepatosteatosis was suppressed by the loss of *Atg7* (Fig. 1C and D). Consistent with the morphological analyses, the amount of TG in control livers increased upon fasting, but such increase was milder in mutant livers (Fig. 1E). Similarly, fasting-dependent hepatosteatosis was blocked in livers of *Atg5*^*f/f*^;Mx1-*Cre* mice one to two weeks after intraperitoneal injection of polyinosinic-polycytidylic acid (pIpC), which induced liver-specific deletion of *Atg5*, another gene essential for autophagy (Supplementary Fig. S1).

### Impairment of partial hepatectomy-induced hepatosteatosis in liver-specific *Atg7*-knockout mice

After partial (70%) hepatectomy, the remnant liver recovers to its original liver weight within approximately one week following hypertrophy of hepatocytes and about two rounds of cell division (Taub, 2004). The liver shows a transient and prominent accumulation of fatty acids one day after resection (Schofield, Sugden, Corstorphine, & Zammit, 1987; Tijburg, Nyathi, Meijer, & Geelen, 1991), which supports rapid cell division and tissue regrowth (Shteyer, Liao, Muglia, Hruz, & Rudnick, 2004). Next, we investigated whether autophagy is also involved in LD biogenesis in hepatocytes after hepatectomy. To this end, we carried out a 70% hepatectomy on the livers of *Atg7*^*f/f*^ and *Atg7*^*f/f*^;Alb-*Cre* mice and followed them until 168 h after hepatectomy. The blood level of free fatty acids in control *Atg7*^*f/f*^ mice gradually decreased after 70% hepatectomy, was at the lowest level at 18–24 h, and recovered 96–168 h after the hepatectomy (Fig. 2A). In contrast, such fluctuation was not observed in mutant *Atg7*^*f/f*^;Alb-*Cre* mice (Fig. 2A), suggesting the impairment of free fatty acid uptake from blood in mutant hepatocytes.

**Figure 2.**
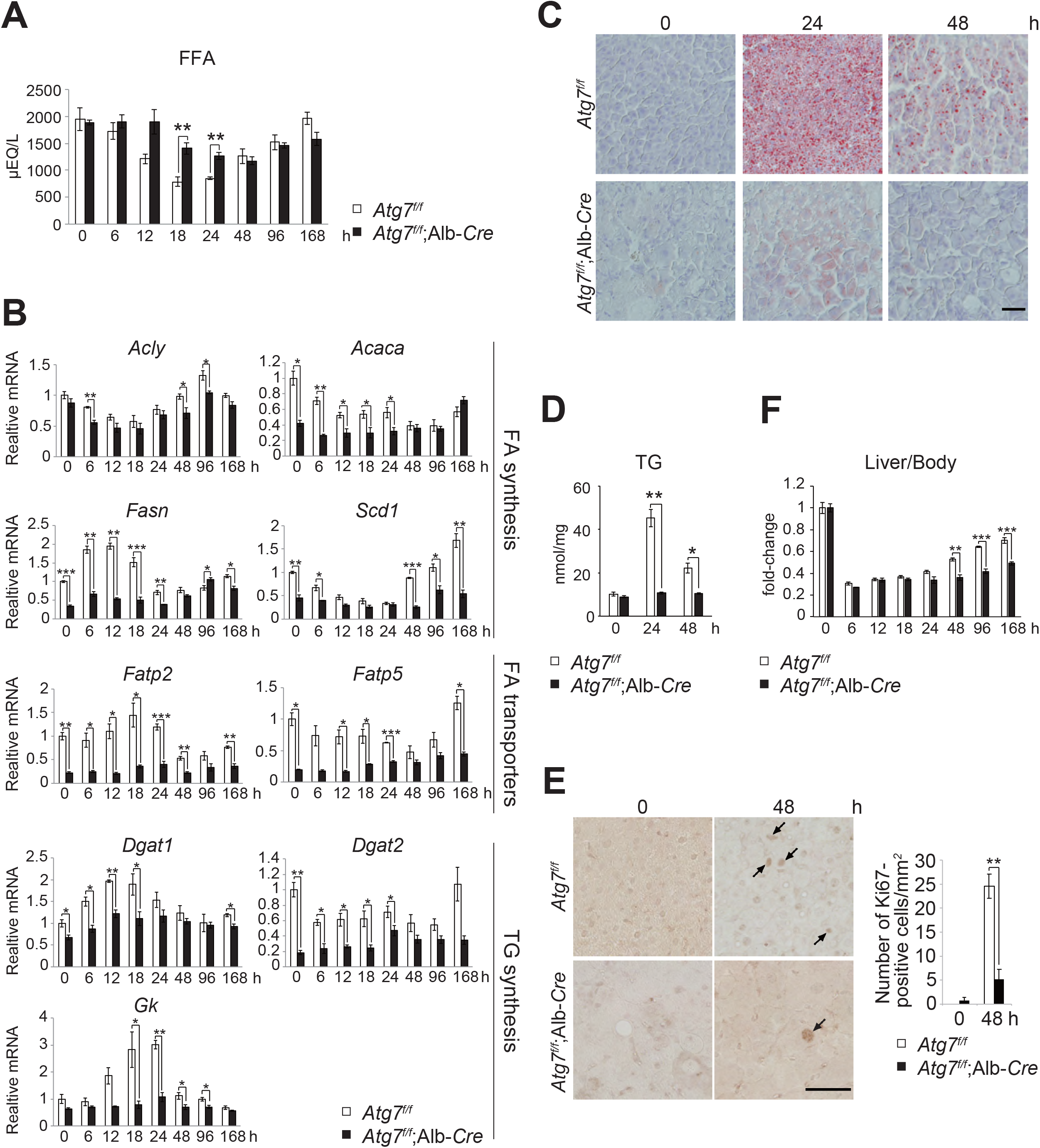
(**A**) Blood-free fatty acid. five-week-old *Atg7*^*f/f*^ (n = 4) and *Atg7*^*f/f*^;Alb-*Cre* (n = 3) mice were subjected to 70% hepatectomy. Subsequently, blood-free fatty acid was measured at the indicated time points following hepatectomy. Data are means ± s.e. \*\**p* < 0.01 as determined by Welch’s *t*-test. (**B**) Gene expression of proteins related to FA and TG syntheses in *Atg7*-deficient livers after partial hepatectomy. Total RNAs were prepared from mouse livers described in (A). Values were normalized against the amount of mRNA in *Atg7*^*f/f*^ livers immediately after the hepatectomy. The experiments were performed three times. Data are means ± s.e. \**p* < 0.05, \*\**p* < 0.01, and \*\*\**p* < 0.001 as determined by Welch’s *t*-test. (**C**) Oil-red O staining. five-week-old *Atg7*^*f/f*^ and *Atg7*^*f/f*^;Alb-*Cre* mice were subjected to 70% hepatectomy. Liver sections were prepared at 0, 24, and 48 h after the hepatectomy and stained with Oil-red O. Data are representative of three separate experiments. Bar: 50 μm. (**D**) Liver triglyceride (TG) in mice described in (A). Data are means ± s.e. \**p* < 0.05, and \*\**p* < 0.01 as determined by Welch’s *t*-test. (**E**) Ki67-staining in liver paraffin sections prepared from five-week-old *Atg7*^*f/f*^ and *Atg7*^*f/f*^;Alb-*Cre* mice at 0 and 48 h following hepatectomy. Number of Ki-67-positive cells (arrows) were counted and plotted as the number per mm^2^ in the right graph (n = 3 mice). Bars: 50 μm. Data are means ± s.e.m. \**p* < 0.05 and \*\*\**p* < 0.001, as determined by Welch’s *t*-test. (**F**) Liver weights (% per body weight) of mice described in (A). Data are means ± s.e. \*\**p*_*vov*_ < 0.01, and \*\*\**p* < 0.001 as determined by Welch’s *t*-test.

While the expression of the FA transporter genes, *Fatp2* and *Fatp5*, in control livers was maintained up to 24 h after the hepatectomy, their expression levels in mutant livers were markedly decreased throughout the time course (Fig. 2B). Moreover, we observed that in control livers, the transcription of genes that encode rate-limiting enzymes related to TG synthesis such as *Dgat1* and *Gk* was dramatically increased up to 24 h. This induction was suddenly terminated 48 h after hepatectomy (Fig. 2B). In contrast, the expression of enzymes involved in FA synthesis dropped to its lowest level 24 h and only recovered to or exceeded the basal level 96–168 h after hepatectomy (Fig. 2B). In mutant livers, expression of almost all genes involved in both FA and TG synthesis, except *Acly*, were suppressed, especially during the early recovery phase following hepatectomy (Fig. 2B). These results suggest that steatosis is defective in autophagy-deficient livers.

Oil-red O staining indicated hyperaccumulation of LDs in control hepatocytes 24 h after hepatectomy and near recovery 48 h after hepatectomy (Fig. 2C). In contrast, such accumulation of LDs was not detectable in *Atg7*-deficient hepatocytes (Fig. 2C). Consistent with those results, in control livers, the amount of TG markedly increased 24 h after hepatectomy and decreased 48 h after hepatectomy (Fig. 2D). Such fluctuation was not observed in the case of mutant livers (Fig. 2D). The hepatosteatosis that follows hepatectomy has been shown to play an essential role in hepatocyte proliferation (Shteyer et al., 2004). Therefore, we speculated that the loss of *Atg7* is accompanied by the impairment of liver regeneration after partial hepatectomy. Indeed, immunohistochemical analysis with anti-Ki67 antibody showed fewer Ki67-positive cells in mutant livers compared with control livers (Fig. 2E). Liver weight per unit body weight in control mice indicated 80% recovery after 168 h compared to the weight before hepatectomy (Fig. 2F). Recovery was observed even in mutant mice, but to a much lesser extent (Fig. 2F).

### NCoR1 degradation through autophagy is necessary for LD biogenesis

Ultimately, we sought to elucidate the molecular mechanism by which loss of autophagy impairs LD formation and utilized HepG2 cells lacking *ATG7* (Supplementary Fig. S2). We expressed either *lacZ* or *ATG7* in *ATG7*-knockout HepG2 cells (#14) and compared the resulting phenotypic differences. As shown in Figure 3A, the conversion of LC3-I to LC3-II was restored in *ATG7^−/−^*HepG2 cells by the expression of *ATG7* but not *lacZ*, and the level of p62 protein decreased upon overexpression of *ATG7* but not *lacZ*. These results support the idea that autophagy is restored in *ATG7^−/−^*HepG2 cells expressing *ATG7* (autophagy-competent) but not in cells expressing *lacZ* (autophagy-incompetent). Indeed, both the nuclear and cytoplasmic NCoR1 levels were lower in autophagy-competent HepG2 cells than in incompetent cells (Fig. 3A). In contrast, we observed a higher level of nuclear LXRα protein in autophagy-competent cells (Fig. 3A). The expression of LXRα target genes in autophagy-competent cells was much higher than in incompetent cells (Fig. 3B). BODIPY-staining revealed that LDs still form in autophagy-incompetent cells (Fig. 3C), but the number and size of LDs were significantly smaller than those in autophagy-competent cells (Fig. 3C). In agreement with these results, we found that the amount of TG in autophagy-competent cells was higher than in incompetent cells (Fig. 3D).

**Figure 3.**
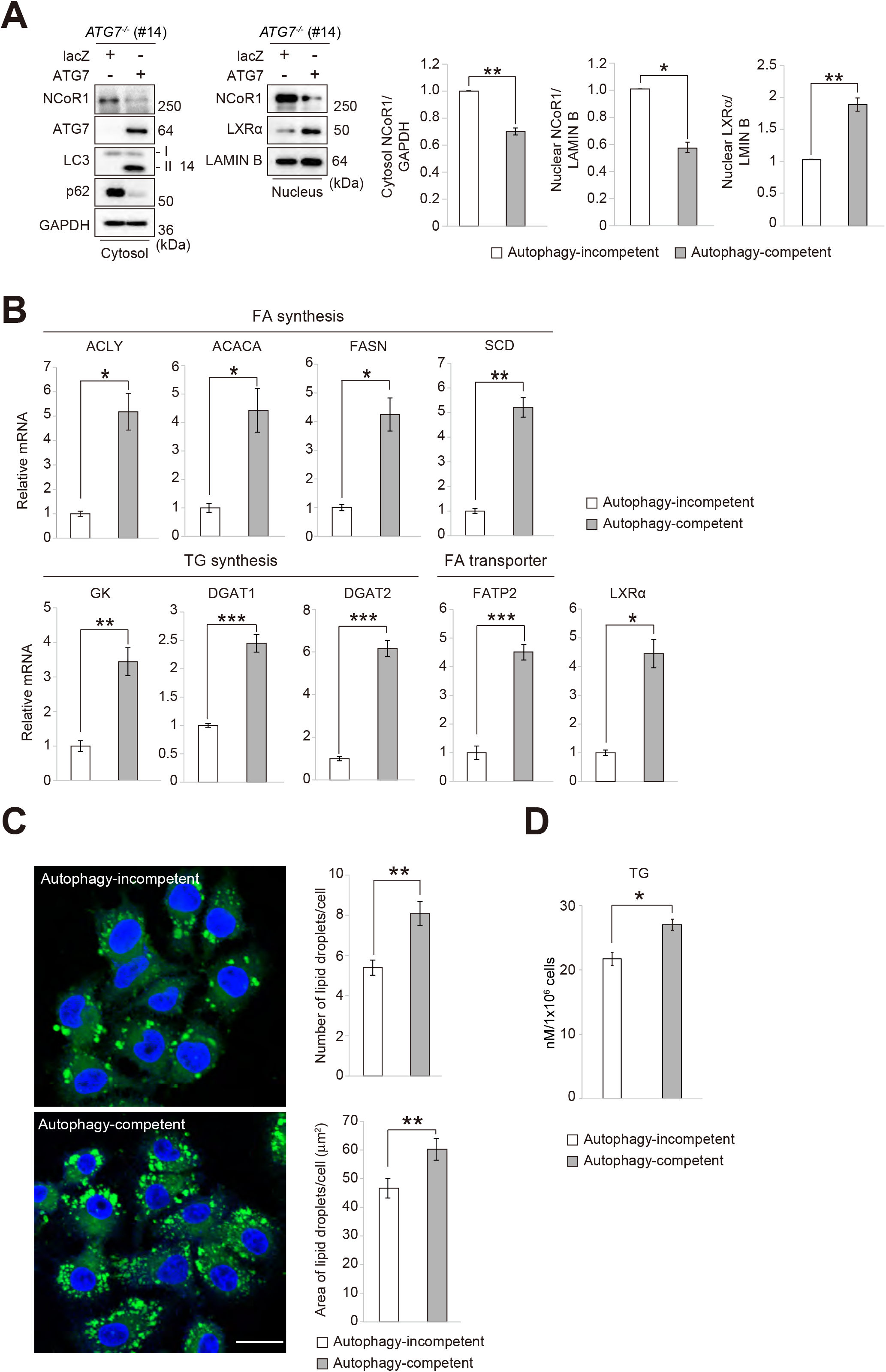
(**A**) Immunoblot analysis. Either lacZ or ATG7 was expressed in *ATG7*-knockout HepG2 (#14) cells using an adenovirus system. Ninety-six hours after infection, both nuclear and cytoplasmic fractions were prepared and subjected to immunoblotting with the indicated antibodies. Data shown are representative of three separate experiments. Bar graphs indicate the quantitative densitometric analyses of nuclear NCoR1 and LXRα relative to LAMIN B and cytoplasmic NCoR1 relative to GAPDH. Statistical analyses were performed using Welch’s *t*-test. Data are means ± s.e. \**p* < 0.05, and \*\**p* < 0.01 as determined by Welch’s *t*-test. (**B**) RT qPCR analysis. Total RNAs were prepared from cells described in (A). Values were normalized against the amount of mRNA in *ATG7*-knockout HepG2 cells (#14) expressing *lacZ* (Autophagy-incompetent). The experiments were performed three times. Data are means ± s.e. \**p* < 0.05, and \*\**p* < 0.01 as determined by Welch’s *t*-test. (**C**) BODIPY staining. Either lacZ or ATG7 was expressed in *ATG7*-knockout HepG2 (#14) cells using an adenovirus system. Ninety-six hours after infection, the cells were stained by BODIPY. Bars: 20 μm. The number and size of LDs were quantified by CellInsight™ CX5 High-Content Screening (HCS) Platform. Statistical analyses were performed using Welch’s *t*-test. Data are means ± s.e. \*\**p* < 0.01 as determined by Welch’s *t*-test. (**D**) Triglyceride levels. Lysates were prepared from cells described in (A), and the concentration of TG in each lysate was determined by using the Abcam Triglyceride Assay Kit. Statistical analysis was performed using Welch’s *t*-test. Data are means ± s.e. \**p* < 0.05 as determined by Welch’s *t*-test.

Next, we investigated whether NCoR1 accumulation in autophagy-incompetent cells directly affects the formation of LDs. The reduced level of nuclear LXRα in autophagy-incompetent cells was increased by the knockdown of *NCoR1* (Fig. 4A). *NCoR1* depletion restores gene expression of most LXRα targets in autophagy-incompetent cells (Fig. 4B). Unexpectedly, the transcription of some LXRα targets in autophagy-incompetent cells, including *Fatp2* and *Dgat1*, did not increase following *NCoR1* ablation (Fig. 4B), probably due to partial compensation by NCoR2, an NCoR1 family protein (Shimizu et al., 2015). *NCoR1* knockdown in autophagy-incompetent cells had little effect on the size and number of LDs (Fig. 4C) since *NCoR1* ablation enhances both the anabolism and catabolism of FAs (Mottis et al., 2013; Perissi et al., 2010). Regardless, we confirmed that the amount of TG in autophagy-incompetent cells was restored to a significant extent by silencing *NCoR1* (Fig. 4D). On balance, we conclude that NCoR1 accumulation due to defective autophagy suppresses LXRα transactivation, resulting in the impairment of FA and TG syntheses and of LD formation.

**Figure 4.**
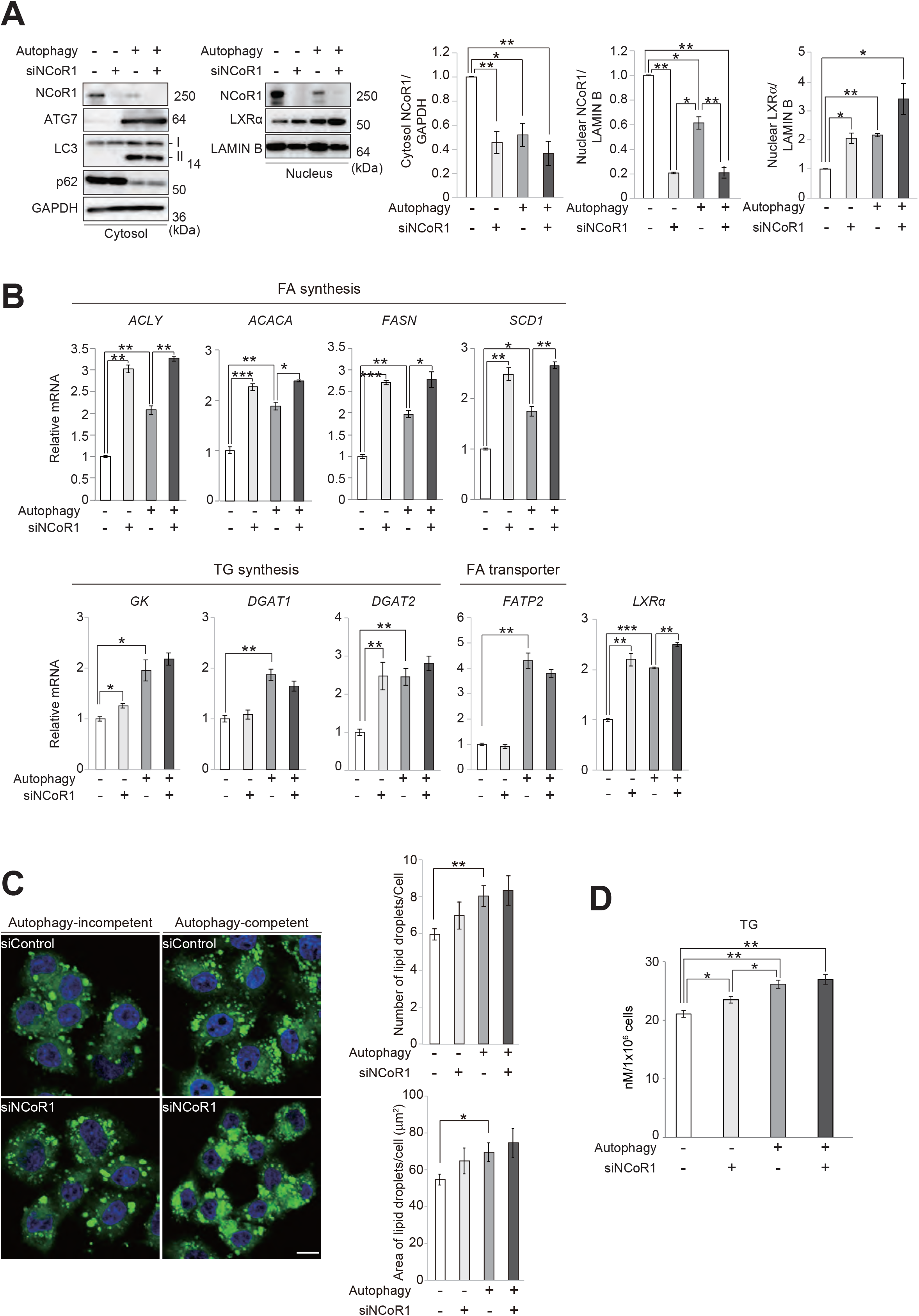
(**A**) Immunoblot analysis. *ATG7*-knockout HepG2 (#14) cells were infected with adenovirus for lacZ (Autophagy -) or ATG7 (Autophagy +) expression. The cells were simultaneously treated with either *NCoR1*-specific or scrambled control siRNA. Forty eight hours after infection, the cells were re-treated with either *NCoR1*-specific or control siRNA and cultured for an additional 48 h. Thereafter, both nuclear and cytoplasmic fractions were prepared and subjected to immunoblotting with the indicated antibodies. Data are representative of three separate experiments. Bar graphs indicate the quantitative densitometric analyses of nuclear NCoR1 and LXRα relative to LAMIN B and cytoplasmic NCoR1 relative to GAPDH. Statistical analyses were performed using Welch’s *t*-test. Data are means ± s.e. \**p* < 0.05, and \*\**p* < 0.01 as determined by Welch’s *t*-test. (**B**) RT qPCR analysis. Total RNAs were prepared from cells described in (A). Values were normalized against the amount of mRNA in *lacZ*-expressing *ATG7*-knockout HepG2 cells treated with control siRNA. The experiments were performed three times. Data are means ± s.e. \**p* < 0.05, \*\**p* < 0.01, and \*\*\**p* < 0.001 as determined by Welch’s *t*-test. (**C**) BODIPY staining. *ATG7*-knockout HepG2 (#14) cells were treated as described in (A). Bars: 20 μm. The number and size of LDs were quantified by CellInsight™ CX5 High-Content Screening (HCS) Platform. Statistical analyses were performed using Welch’s *t*-test. Data are means ± s.e. (**D**) Triglyceride levels. Lysates were prepared from cells described in (A), and the concentration of TG in each lysate was determined using the Abcam Triglyceride Assay Kit. Statistical analysis was performed using Welch’s *t*-test. Data are means ± s.e. \**p* < 0.05, and \*\**p* < 0.01 as determined by Welch’s *t*-test.

Our finding differs from a prior report that mouse hepatocytes lacking *Atg7* increase the size and the number of LDs (Singh et al., 2009). The main difference in experimental settings between this prior study and ours is the age of the genetically modified *Atg7*^*f/f*^; Alb-*Cre* mice. While we used five-week old *Atg7*^*f/f*^; Alb-*Cre* mice, Singh R *et al.*, utilized four-month old mice (Singh et al., 2009). Under conditions where the regeneration of mature hepatocytes is defective, such as the lack of β-catenin, hepatic oval cells proliferate and differentiate into hepatocytes and cholangiocytes, both of which replace the liver mass with aging (Wang et al., 2011). During active proliferation, most hepatic progenitor cells derived from oval cells undergo maturation arrest and become dedifferentiated, but these progenitor-derived immature hepatocytes possess a high potential for developing into liver tumors. Hepatocyte-specific ablation of *β-catenin*, in fact, promotes tumorigenesis (Wang et al., 2011). Likewise, the long-term suppression of autophagy in mouse livers is accompanied by tumorigenesis (Inami et al., 2011; Takamura et al., 2011), suggesting that autophagy-defective hepatocytes may lose the ability to regenerate and that the hepatic oval cells may compensate it. Remarkably, both hepatocytes and cholangiocytes differentiated from oval cells express albumin at negligible levels (Wang et al., 2011). In fact, we observed the presence of hepatocytes that lack p62-positive structures mosaically in aged *Atg7*^*f/f*^; Alb-*Cre* mice (*i.e.*, autophagy-competent hepatocytes) (data not shown). Thus, we speculate that the livers of aged *Atg7*^*f/f*^; Alb-*Cre* mice are partially composed of Atg7-intact hepatocytes derived from oval cells and that such hepatocytes accumulate LDs aggressively to compensate for the dysfunction of *Atg7*-deficient hepatocytes.

Autophagy provides a substantial amount of FAs through the degradation of organelles under nutrient-deprived conditions (Rambold et al., 2015). A robust influx of FAs from the blood into peripheral cells such as hepatocytes occurs under fasting conditions or after hepatectomy (Schofield et al., 1987; Tijburg et al., 1991). The increased level of intracellular FAs due to starvation-induced autophagy and/or fasting-triggered influx provides fuel for β-oxidation to produce energy; however, since β-oxidation intermediates such as acylcarnitine have a cytotoxic effect (Nguyen et al., 2017), cells have to maintain the levels of intracellular FAs below certain limits. In fact, DGAT1-mediated TG synthesis under nutrient-deprived conditions, when the amount of intracellular FAs is excessive, is necessary to mitigate the lipotoxic cellular damage caused by acylcarnitine (Nguyen et al., 2017). It is worth noting that the loss of autophagy in mouse livers is accompanied by NCoR1 accumulation, resulting in increased levels of acylcarnitine, in particular under fasting conditions (Iershov et al., 2019; Saito et al., 2019). We conclude that fine-tuning of NCoR1 protein levels through autophagy regulates LD formation in order to mitigate lipotoxicity.

## Materials and methods

### Cell culture

HepG2 cells were grown in Dulbecco’s modified Eagle medium (DMEM) containing 10% fetal bovine serum (FBS), 5 U/ml penicillin, and 50 μg/ml streptomycin. For knockdown experiments, HepG2 cells were transfected with 25 nM SMARTpool siRNA for *NCoR1* using Dharmafect 1 (Thermo Fisher Scientific, Waltham, MA, USA). *ATG7*-knockout HepG2 cells (Saito et al., 2019) were used in this study.

### Mice

*Atg7*^*f/f*^ (Komatsu et al., 2005), *Atg7*^*f/f*^;Alb-*Cre* (Komatsu et al., 2007), *Atg5*^*f/f*^ (Hara et al., 2006), and *Atg5*^*f/f*^;Mx1-*Cre* (Hara et al., 2006) mice in the C57BL/6 genetic background were used in this study. Mice were housed in specific pathogen–free facilities, and the Ethics Review Committees for Animal Experimentation of Niigata University, the University of Tokyo, and Juntendo University approved the experimental protocol. The concentration of liver triglycerides was determined using the Triglyceride Assay Kit, ab65336 (Abcam, Cambridge, UK). Free fatty acids in plasma were analyzed by SRL (Tokyo, Japan). Partial hepatectomy (PHx) was performed in six-week-old male mice. Mice were anesthetized with an intraperitoneal injection of 0.05 ml/10 g body weight of a mixed anesthetic agents, consisting of medetomidine (0.06 mg/ml), midazolam (0.8 mg/ml), and butorphanol (1 mg/ml) in sterile normal saline and subjected to approximately 70% PHx by removing the left lateral and median lobes after midventral laparotomy. The mortality rate after 70% PHx was <1%.

### Immunoblot analysis

Livers were homogenized in 0.25 M sucrose, 10 mM 2-[4-(2-hydroxyethyl)-1-piperazinyl]ethanesulfonic acid (HEPES) (pH 7.4), and 1 mM dithiothreitol (DTT). Nuclear and cytoplasmic fractions from livers and cultured cells were prepared using the NE-PER Nuclear and Cytoplasmic Extraction Reagents (Thermo Fisher Scientific). Samples were subjected to SDS-PAGE, and transferred to a polyvinylidene difluoride membrane thereafter (Merck, IPVH00010). Antibodies against LXRα (ab28478, Abcam, Cambridge, UK; 1:500), PPARα (ab8934, Abcam, Cambridge, UK; 1:500), NCoR1 (#5948S, Cell Signaling Technology, Danvers, MA, USA; 1:500), Atg7 (013-22831, Wako Pure Chemical Industries, Osaka, Japan; 1:1000), p62 (GP62-C, Progen Biotechnik GmbH, Heidelberg, Germany; 1:1000), LC3B (#2775, Cell Signaling Technology; 1:500), Gapdh (MAB374, Merck Millipore Headquarters, Billerica, MA, USA; 1: 1000), and lamin B (M-20, Santa Cruz Biotechnology; 1:200) were purchased from the indicated suppliers. Blots were incubated with horseradish peroxidase-conjugated goat anti-mouse IgG (H+L) (Jackson ImmunoResearch Laboratories, Inc., 115-035-166), goat anti-rabbit IgG (H+L) (Jackson ImmunoResearch Laboratories, Inc., 111-035-144) or goat anti-guinea pig IgG (H+L) antibody (Jackson ImmunoResearch Laboratories, Inc., 106-035-003), and visualized by chemiluminescence. Band density was measured using the software Multi Gauge V3.2 (FUJIFILM Corporation, Tokyo, Japan).

### RT-qPCR (Real-time quantitative reverse transcriptase PCR)

Using the Transcriptor First-Strand cDNA Synthesis Kit (Roche Applied Science, Indianapolis, IN, USA), cDNA was synthesized from 1 μg of total RNA. RT qPCR was performed using the LightCycler^®^ 480 Probes Master mix (Roche Applied Science) on a LightCycler^®^ 480 (Roche Applied Science). Signals from human and mouse samples were normalized against *GAPDH* (glyceraldehyde-3-phosphate dehydrogenase) and *Gusb* (ß-glucuronidase) mRNA, respectively. The sequences of primers used for gene expression analysis in either mouse livers or human cell lines are provided in Supplementary Table S1.

### Histological examinations

Excised liver tissues were fixed by immersing in 0.1 M phosphate buffer (PB, pH 7.4) containing 4% paraformaldehyde and 4% sucrose. They were embedded in frozen OCT-compound or paraffin. The cryo-sections were stained with Oil-red O, and paraffin sections were stained with rabbit anti-Ki67 antibody (clone SP6, Thermo Fisher Scientific) followed by N-Histofine^®^ simple stain mouse MAX PO kit (NICHIREI BIOSCIENCES, Japan) using 3,3’-diaminobenzidine. They were observed with a light microscope (BX51, Olympus). For quantification, Ki67-positive cells were counted in 5 rectangular regions (433 x 326 ◻m) per liver section of each mouse. Three to four mice were included in this analysis.

### Electron microscopy

Livers were fixed by immersing in 0.1 M PB containing 2% paraformaldehyde and 2% glutaraldehyde. They were post-fixed with 1% OsO_4_, embedded in Epon812, and sectioned for observation with an electron microscopy (JM-1200EX, JEOL, Japan). For quantification, area ratio of LDs was measured in 20 hepatocytes for each mouse. Three mice were included in this analysis.

### Microscopy for cultured cells

For staining of lipid droplets, cells were incubated with 1 μg/ml BODIPY™ 493/503 (D3922, Thermo Fisher Scientific) diluted in PBS for 15 min and extensively washed with PBS. Finally, the cells were incubated for 5 min with 10 μg/ml of Hoechst 33342 diluted in PBS, washed with PBS and mounted on slides with Prolong Gold antifade mounting solution (Thermo Fisher Scientific). Ten fields of cells were imaged for each experimental condition with a CellInsight™ CX5 High-Content Screening (HCS) Platform (Thermo Fisher Scientific) using HCS Studio™ software.

### Statistical analysis

Values, including those displayed in the graphs, are means ± s.e. Statistical analysis was performed using the unpaired t-test (Welch test). A P value less than 0.05 was considered to indicate statistical significance.

## Acknowledgements

We thank K. Kanno (Fukushima Medical University) for his help in histological analyses. M.K. is supported by Grants-in-Aid for Scientific Research on Innovative Areas (19H05706 to M.K), the Japan Society for the Promotion of Science (an A3 foresight program, to M.K. and 18H02611 to M.K.), and the Takeda Science Foundation (to M.K.).

---

**Supplementary Figure S1.**
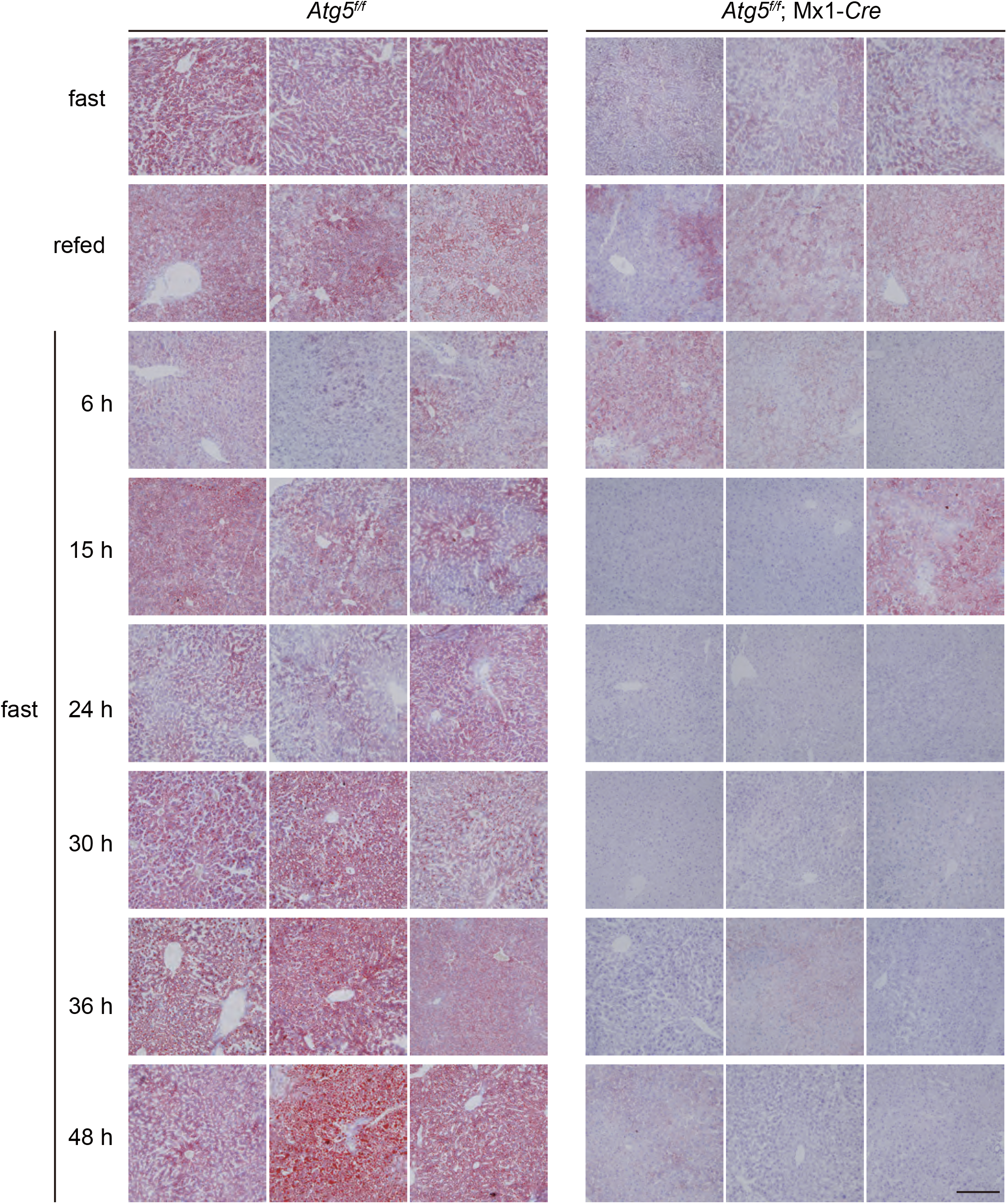
Representative Oil-red O staining images of hepatocytes of 12-week-old *Atg5*^*f/f*^ and *Atg5*^*f/f*^;Mx1-*Cre* mice. *Atg5*^*f/f*^ and *Atg5*^*f/f*^;Mx1-*Cre* mice were injected intraperitoneally with pIpC to delete *Atg5* in the liver at 10 weeks of age. Bars: 200 μm.

**Supplementary Figure S2.**
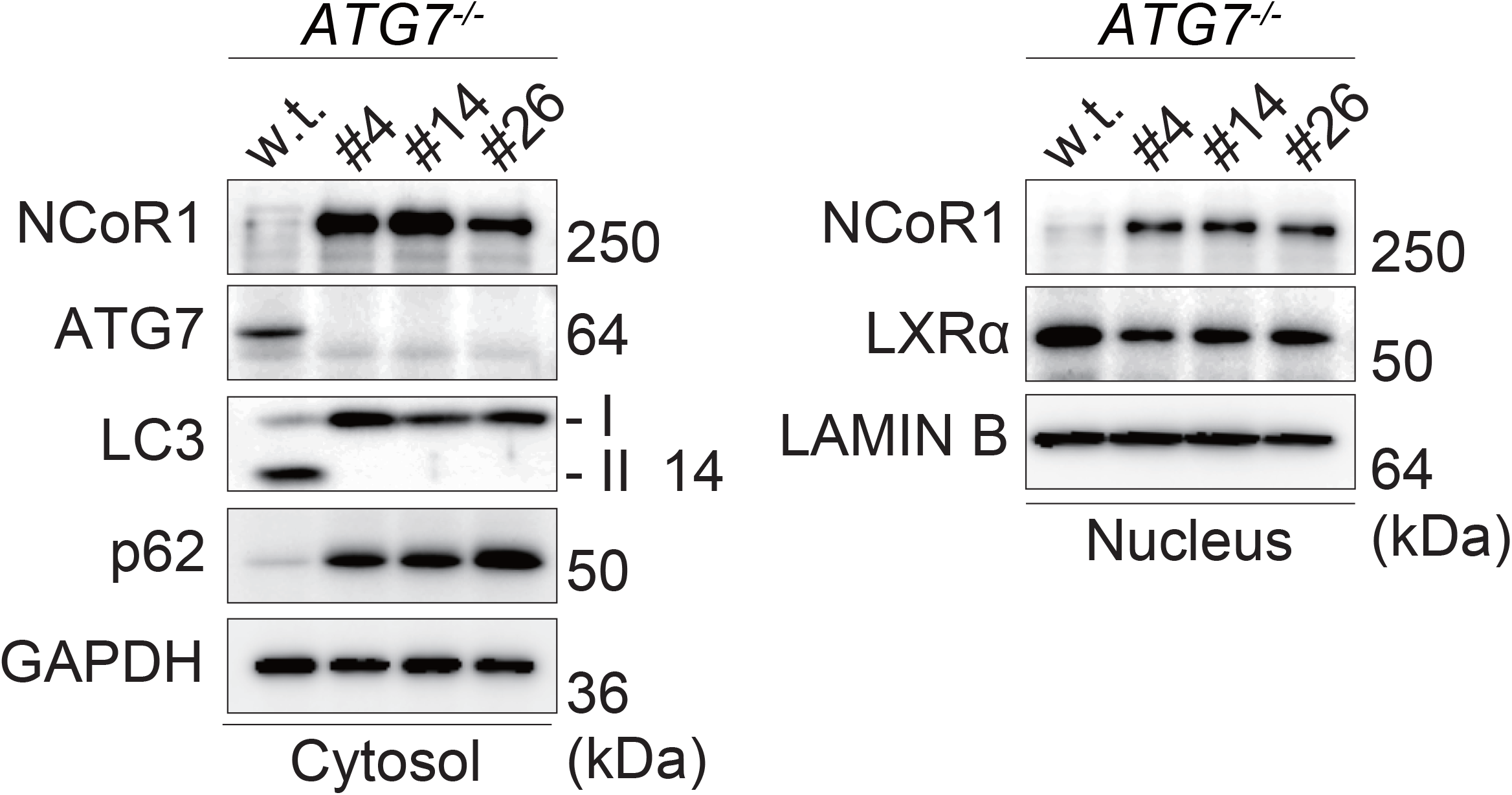
Immunoblot analysis. Both nuclear and cytoplasmic fractions were prepared from parental and *ATG7*-knockout HepG2 cells (#4, #14, and #26) and subjected to immunoblotting with the indicated antibodies. Data are representative of three separate experiments.

**Supplementary Table 1.**
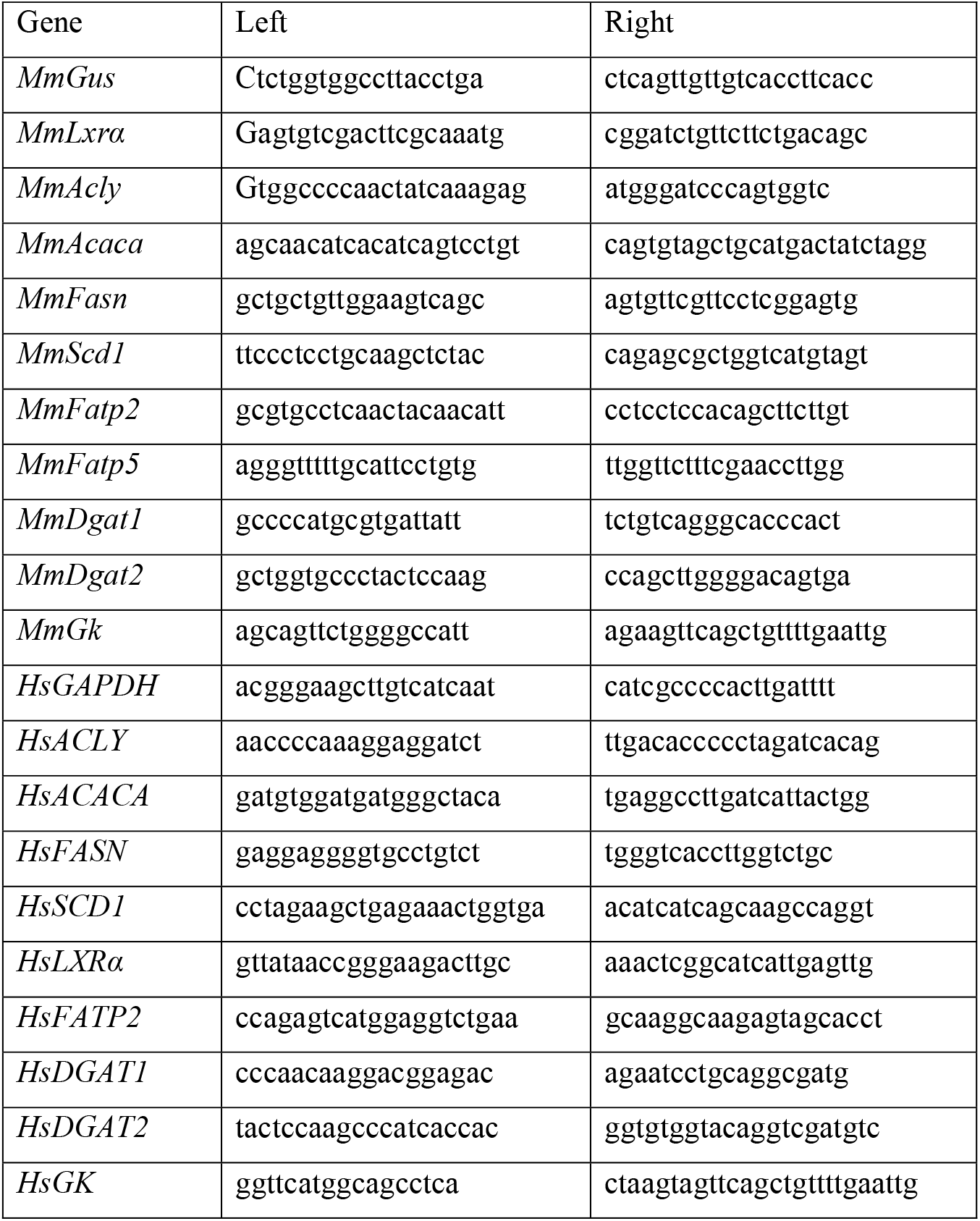
Primer sequences used for RT qPCR.

